# Structural enzymological studies of the long chain fatty acyl-CoA synthetase FadD5 from the *mce1* operon of *Mycobacterium tuberculosis*

**DOI:** 10.1101/2024.10.29.620959

**Authors:** Mohammad Asadur Rahman, Subhadra Dalwani, Rajaram Venkatesan

## Abstract

The cell wall of *Mycobacterium tuberculosis* (Mtb), the causative agent of tuberculosis, is rich in complex lipids. During intracellular stage, Mtb depends on lipids for its survival. Mammalian cell-entry (Mce) 1 complex encoded by the *mce1* operon is a mycolic/fatty acid importer. *mce1* operon also encodes a putative fatty acyl-CoA synthetase (FadD5; Rv0166), potentially responsible for the activation of fatty acids imported through the Mce1 complex by conjugating them to Coenzyme A. Here we report that FadD5 is associated to membrane although it can be purified as a soluble dimeric protein. ATP and CoA binding influence FadD5’s stability and conformation respectively. Enzymatic studies with fatty acids of varying chain lengths show that FadD5 prefers long chain fatty acids as substrates. X-ray crystallographic studies on FadD5 and its variant reveal that the C-terminal domain (∼100 residues) is cleaved off during crystallization. Noteworthy, deletion of this domain renders FadD5 completely inactive. SAXS studies, however, confirm the presence of full length FadD5 as a dimer in solution. Further structural analysis and comparisons with homologs provide insights on the possible mode of membrane association and fatty acyl tail binding.

## 1. Introduction

Tuberculosis (TB) is an alarming global health issue and millions of new cases are being reported every year with more than a million deaths every year [1]. *Mycobacterium tuberculosis* (Mtb), the causative agent of TB, demonstrates remarkable metabolic versatility due to its unique fatty acid metabolism system. Mtb possesses a large number of genes involved in fatty acid metabolism. Among the lipid metabolizing genes, 34 genes are designated as *fadD*s [2] which code for acyl activating enzymes (AAE). Sequence alignment and a phylogenetic tree, show two subgroups of FadDs, namely, fatty acyl-AMP ligase (FAAL) (EC 6.2.1.20) and the fatty acyl-CoA synthetase (FACSs) (EC 6.2.1.3) [3]. FAALs activate fatty acids by converting them to acyl-adenylates and transfer them to modular enzymes such as polyketide synthases for chain extension or biosynthesis of lipidic metabolites. Most frequently, genes coding for FAAL are present adjacent to genes coding for polyketide synthases [4–6]. FACSs, on the other hand, activate fatty acids by conjugating them to coenzyme A (CoA) [7]. The addition of CoA to the fatty acids is the first step needed for further processing of fatty acids by other pathways such as β-oxidation. FACS converts fatty acids to fatty acyl-CoAs in a two-step reaction. One distinguishing difference from FAAL to FACS is the presence of ∼20 amino acid insertion motif in FAAL [4]. FadDs contain the conserved AAE fold with two domain structure with a large N-terminal domain and a smaller C-terminal domain. The substrate binding pocket is located between the two domains [4]. In FACS, during catalysis, C-terminal domain moves from adenylate to thiolation conformation which is known as domain alternation [8,9]. One of the key characteristics of long chain FACS enzymes is their membrane bound nature which is important for anchoring large substrates [10,11]. Structural studies have been conducted for FadDs from Mtb, such as FadD10, FadD13, FadD23, FadD28 and FadD32 in the unliganded form or in the presence of different ligands and in some cases also with only the N-terminal domain [3,7,11–13].

Function of *mce1* operon has been studied extensively and the structural studies have been conducted on individual Mce proteins as well as full Mce1 complex [14–17]. *mce1* operon has a gene *fadD5*, a putative FACS which is co-transcribed with the other genes in the *mce1* operon [18]. Previous studies showed that mice infected with *fadD5* mutant survived significantly longer than those infected with the wild type Mtb [19] suggesting that FadD5 is a potential drug target. This study also showed that *fadD5* mutant grows slower in minimal media supplemented with mycolic acid compared to other long chain fatty acid as a sole carbon source [19] suggesting FadD5 being involved in mycolic acid recycling during infection. Moreover, *mce1* operon mutant also shows an accumulation of mycolic acid in its cell wall which also suggests that Mce1 complex is a mycolic acid ABC transporter [15]. Whenever *E.coli fadD* was complemented with Mtb *fadD*5, it was unable to utilize long chain fatty acid as a sole carbon source, demonstrating the uniqueness of FadD5 which indicate its different function or substrate specificity [19]. Though FadD5 has been studied *in vivo* no structural and enzymatic studies have been carried out so far.

Considering the importance of FadD5 involved in activating the lipids transported by the Mce1 complex, this study is about the structural and functional characterization of FadD5. Biophysical studies suggested that although FadD5 is stable as a dimeric enzyme in solution, it is also associated to membrane. Interestingly, the C-terminal domain was cleaved off during crystallization. Enzymatic studies demonstrated its FACS activity with chain length of up to 24. Structural and modelling studies provide insights on the possible mode of fatty acyl tail binding. Overall, the studies provide insight on the structural and enzymatic properties of FadD5, an important enzyme for the utilization/recycling of lipids imported by the Mce1 complex.

## 2. Materials and methods

### 2.1 Sequence analysis

The sequence similarity was analyzed by the Basic Local Alignment Search Tool (BLAST) program from the NCBI, USA server against protein data bank [20]. MycoBrowser portal (http://mycobrowser.epfl.ch) was used as a source to get gene and protein information of Mtb [21]. Multiple sequence alignment was performed by the T-Coffee program on EMBL-EBI website [22] and sequence alignment was visualized with ESPript3 [23].

### 2.2 Cloning and expression of fadD5

*fadD5* was PCR amplified and cloned into MCS-1 of pETDuet-1 vector using a restriction-based cloning approach using EcoRI and HindIII restriction enzymes resulting in a clone coding for His_6_ tag at the N-terminus. The genomic DNA of Mtb H37Rv (purchased from ATCC) was used as the template. A deletion construct coding for FadD5_1-448_ was prepared, truncating the region coding for the C-terminal domain. All the constructs made were confirmed by DNA sequencing. The constructed vectors pETDuet-*fadD5* and pETDuet-*fadD5*_1-448_ were transformed into chemically competent *E. coli* BL21 cells. Overnight-grown cultures were added into new LB media and grown at 30 °C until the OD_600_ reached 0.6 and expression of *fadD5* was induced by adding 0.1 mM isopropyl β-D-thiogalactopyranoside (IPTG) at 20 °C.

### 2.3 Purification of FadD5

The cells expressing FadD5 and FadD5_1-448_ were harvested by centrifugation at 4000 g for 20 mins at 4 °C. The harvested bacterial pellet was resuspended in lysis buffer (20 mM HEPES pH 7.5, 500 mM NaCl, 5 mM MgCl_2_, 5 mM Imidazole, 1 mM β-mercaptoethanol (BME)) with one EDTA-free protease inhibitor tablet, lysozyme, DNase and RNase. The cells were lysed using a French press and the soluble fraction was collected by centrifuging for 40 min at 16000 g at 4 °C. Then, FadD5 was purified by immobilized metal affinity chromatography (IMAC) by washing with the lysis buffer and eluted with a buffer containing 400 mM imidazole. The eluted protein was concentrated and injected into Superdex 200 10/300 (Cytiva) size-exclusion chromatography (SEC) column pre-equilibrated with 20 mM HEPES buffer containing 500 mM NaCl, 5 mM MgCl_2_, 1 mM BME at pH 7.5. The peak fractions were collected, concentrated, flash-frozen in aliquots and stored at −70 °C for further use. The fractions were analyzed by 12% SDS-PAGE and the identity of the protein was confirmed by peptide mass finger printing analysis.

### 2.4 Membrane association studies of FadD5

For membrane association studies, FadD5 and FadD5_1-448_ expressed cells were lysed using a French press and cell debris were removed by centrifugation at 6000 g. From the resulting supernatant, membrane fraction was isolated by ultracentrifugation at 45000 g for 1 h at 4 °C. The pellet was washed twice by resuspending in a buffer containing 20 mM HEPES, 500 mM NaCl, 5 mM MgCl_2_, 1 mM BME at pH 7.5 and membrane fraction was isolated by centrifugation again. The membrane was solubilized in the membrane extraction buffer containing 10 mM DDM. FadD5 was purified from the solubilized fraction as mentioned earlier and analyzed by SDS-PAGE and peptide mass fingerprinting analysis.

### 2.5 Blue native PAGE (BN-PAGE)

Approximately 40 µg of protein was loaded onto 4-15% mini-PROTEAN TGX precast gel (BIORAD) and run for 2-3 hours at 20 mA at 4 °C using Tris-glycine buffer as anode buffer and Tris-glycine supplemented with 0.02% Coomassie G-250 as the cathode buffer. Bovine serum albumin (BSA) was used as a control. The gel was then destained overnight in water and the bands were analyzed.

### 2.6 Multi-angle laser light scattering (MALLS)

60 µl (1 mg ml^-1^) of purified FadD5 or its variant was injected into a Superdex 200 Increase 10/300 GL column (Cytiva) re-equilibrated with protein buffer coupled to a Shimadzu HPLC/FPLC system, a refractive index (RI) detector (Wyatt Technologies), a UV detector and a MALLS device (Wyatt Mini DAWN Treos instrument, Wyatt Technologies). The molecular weight of the protein was calculated (UV and RI signals were used as the source of concentration) for the eluted peaks using *ASTRA* 7.0 software (Wyatt Technologies).

### 2.7 Circular dichroism (CD) spectroscopy

Protein was diluted in water to a final protein concentration of 0.1 mg ml^-1^. The buffer was also diluted in a similar way to use as the blank. The final buffer composition was 2 mM HEPES, pH 7.5, 50 mM NaCl, 0.1 mM BME. CD spectroscopy data were collected in a Chirascan CD spectrometer (Applied Photophysics, Leatherhead, UK) between 280 and 190 nm at 22 °C in a 0.1 cm path-length quartz cuvette. *Chirascan Pro-Data Viewer* (Applied Photophysics) and *CDNN* software (http://www.xn--gerald-bhm-lcb.de/download/cdnn) were used to process and deconvolute the collected CD data to calculate the secondary structure composition. The direct CD measurements (θ; mdeg) were converted into mean residue molar ellipticity ([θ]_MR_) by Pro-Data Viewer. Thermal unfolding was recorded between 190 and 280 nm from 22 to 92 °C at a rate of 1 °C per min using a Peltier temperature control TC125 (Quantum Northwest, Liberty Lake, WA). Thermal melting temperature (T_m_) was calculated using *Global3* software (Applied Photophysics) using the global fit analysis protocol by fitting the data to a one-transition model.

### 2.8 Nano differential scanning fluorimetry (nanoDSF)

A label-free fluorescence-based approach was used for thermal stability analysis of FadD5 through nanoDSF, using Prometheus NT.48 (NanoTemper Technologies, Germany). 10 µl of the purified protein alone or mixed with 2 mM ATP or 2 mM CoA was loaded into standard glass capillaries for nanoDSF. The fluorescence signal resulting from the thermal unfolding of the protein was measured at 330 nm and 350 nm (F350/F330) by heating the sample from 20 °C to 90 °C by a thermal ramping rate of 1 °C per min. The fluorescence intensity ratio and its first derivatives were calculated with the PR.ThermalControl (NanoTemper Technologies, Germany). The data was further analyzed with MoltenProt (https://spc.embl-hamburg.de/app/moltenprot).

### 2.9 Enzyme activity assay

FadD5 activity was measured by monitoring the first half of the reaction through the measurement of inorganic phosphate (Pi) produced from the hydrolysis of inorganic pyrophsophate (PPi) resulting from the enzyme catalyzed reaction [24,25]. The amount of Pi was quantified using a colorimetric Malachite Green phosphate assay kit (Sigma) at 630 nm. The standard 100 µl reaction contained 50 mM HEPES, pH 7.5, 5 mM MgCl_2_, 1 mM BME, 0.001% Brij-35, 0.5 mM ATP, 0.5 mM CoA, 0.3 U pyrophosphatase, 100 µM fatty acids. The reaction was started by adding 0.5 µg of purified protein and continued for 30 min at RT. The reaction was stopped by incubating the reaction mix at 80 °C in a water bath for 5 min and was diluted with 400 µl of 50 mM HEPES, pH 7.5 buffer. Then 80 µl of the sample was mixed with 20 µl of color developing reagent according to the manufacturer’s recommendation and incubated for 30 min for color development and the absorbance was recorded in a Tecan Spark plate reader (Tecan, Switzerland). Reaction without protein/substrate was used as a blank. The amount of Pi released was calculated from the calibration curve plotted with a known concentration of Pi according to the manufacturer’s instruction. All the experiments were done in triplicates.

### 2.10 Liquid chromatography–mass spectrometry (LCMS) experiment to detect the product formation by FadD5

Standard reaction without CoA was considered as the first half of reaction and addition of CoA was considered as the full reaction. The control reactions were carried out without the enzyme. LCMS was used to detect the formation of C12-AMP and C12-CoA from C12 in a two-step reaction. The standard reaction mix was diluted 500-fold and analysed by LCMS on a Q Exactive system linked to an Aquity UPLC system equipped with ACQUITY UPLC® BEH C18 1.7 µm (2.1 mm x 100 mm) column (Waters Acquity). The column was eluted at 0.3 ml min^-1^ with a gradient of 10 mM ammonium acetate (solvent A) and acetonitrile (solvent B) with the following gradient points: 1 min 99% A, 9 min 10% A, 1 min 99% A. Mass spectra were recorded in negative ion mode from m/z 300 to 1500 at resolution of 70000 for up to 200 msec per scan and automatic gain control target at 3×10^6^. Data were analyzed by the Qual browser option of XCalibur (Thermo Fisher).

### 2.11 Small-angle X-ray scattering (SAXS)

SAXS data were collected at B21 beamline [26] at Diamond Light Source, UK in batch-mode. Measurements were collected for FadD5 at 4 mg ml^-1^ and in the presence of 2 mM ATP or 2 mM CoA, with an appropriate buffer as blank. For each sample, 21 frames with 1 sec exposure per frame were collected. ScÅtter and PRIMUS software [27,28] were used for the initial data processing. The buffer scattering was subtracted from the protein scattering using ScÅtter and the subtracted data was rebinned. The radius of gyration (R_g_), forward scattering I(0), interatomic distance distribution function P(r), maximum particle distance (D_max_) and porod volume (V_p_) were calculated by *ScÅtter* and *PRIMUS*. Molecular weight was calculated using the calculated porod volume and *SAXSmow* [29].

### 2.12 Crystallization studies

FadD5 and its variants were crystalized in TTP plates using the sitting drop vapor diffusion method using an alternative reservoir approach [30] using Mosquito robot (SPT Labtech). MIADSplus commercial screen (Molecular dimensions) was used as the reservoir solution. Freshly purified, FadD5 or its variants were mixed with 5 mM CoA and 1 mM ATP to a final concentration of 18 mg ml^-1^ and mixed with 0.1 M Sodium formate, 20 % w/v SOKALAN CP 45 (crystalizing solution) in 1:3 ratio at room temperature. Crystal appearance was monitored by IceBear [31]. Crystals appear in 1-5 days. The crystalizing solution was mixed with 25% glycerol (v/v) for cryo-protection and flash-frozen in liquid nitrogen for diffraction studies. For binding studies, cryoprotectant solution was supplemented with ATP and CoA.

### 2.13 Data collection, structure determination and structure refinement

X-ray crystallographic data from the crystals of FadD5 and its variants were collected from Diamond Light Source (UK). Data reduction and scaling of the collected data was done by *XDS* [32] and *AIMLESS* [33] or the auto processed data from different data processing software available at Diamond Light Source, such as *fast_dp* [34], *xia 2* [35] were used for further processing. Molecular replacement was done using *Expert MR-PHASER* of *CCP4i2* package [36] using FadD13 structure (PDB id: 3R44) as the search model. The model building was performed with *COOT* [37] and the refinement was done by *REFMAC* from *CCP4i2* [36] or using *Phenix* [38]. The images of the structures were generated, visualized and compared using *PyMOL* (https://www.pymol.org) or *UCSF ChimeraX* [39].

## 3 Results and discussion

### 3.1 Sequence analysis shows the conserved motifs of FadD5

Mtb FadD5 (Uniprot id: O07411, Mycobrowser id: Rv0166) consists of 554 amino acids with a calculated molecular mass of ∼60 kDa. The recombinant MtFadD5 has an additional 16 amino acids with a total calculated molecular mass of ∼62 kDa. Sequence comparison with other FadD homologs shows the presence of the conserved motifs, A1 to A10 and signature motifs in FadD5 as described in previous studies (Fig. S1) [40,41]. Important motifs identified in FadD5 are listed here. i) 27 amino acids long signature motif/FACS motif, responsible for fatty acid binding (^407^GGWFHSGDLVRMDSDGYVWVVDRKKDM^433^). ii) Linker (L) motif responsible for various closed conformations of the C-terminal domain, displays a consensus sequence “Asp-Arg-Xaa-Lys” in all long chain FACSs, while the short chain and medium chain FACSs, the consensus motif sequence is “Gly-Arg-Xaa-Asp” [42,43]. Sequence analysis suggests that FadD5’s L motif is Asp428-Arg429-Xaa-Lys431 (^428^DRKKDM^433^) as observed in other long chain FACS that connect N– and C-terminal domains. iii) P-loop (^187^IMYTSGTTGRPKGA^202^) is responsible for binding phosphate of ATP. iv) A motif (^328^LAAFGQTE^335^) for the binding of adenine ring of ATP/AMP. P-loop and A motif collectively also known as ATP/AMP-binding signature motif [44,45] (Fig. S1). Gate (G) motif (^232^VPLFHIAG^239^) keeps the enzyme in the closed conformation required for catalysis and interact with the fatty acyl chain. G motif plays a role in determining the specific functions, activities and substrate length preferences in FadDs. Firefly luciferase’s G motif contains Phe247 [46], potentially the gating residue for its catalytic activity. Similarly, the gating residue in TtLC-FACS is Trp234 [40]. The gating residue is thought to change conformation to accommodate longer fatty acyl chains. However, interestingly in FadD5, the corresponding residue is less bulky Ile240. Similarly, the gating residue in FadD13 is Leu226 [11] which is also known to bind very long acyl chains.

### 3.2 FadD5 and FadD5_1-448_ are dimer in solution and are peripheral membrane proteins

FACS exists as monomers as well as dimers whereas FAALs mostly function as monomers [11,12,40]. The FadD5 and FadD5_1-448,_ expressed in *E. coli* were purified from the soluble fraction using IMAC followed by SEC which shows that FadD5 and FadD5_1-448_ elute at a retention volume of ∼13.5 ml and ∼14.5 ml respectively, in a 24 ml Superdex 200 10/300 column (Fig. 1A). Blue-native PAGE analysis shows a predominant band of FadD5 close to the dimeric band of BSA corresponding to a molecular mass of 132 kDa suggesting that FadD5 is purified as a homodimer in solution (theoretical molecular mass of 124 kDa) (Fig. 1B). The dimeric nature of FadD5 was further confirmed by SEC-MALLS experiment according to which the calculated molecular mass of FadD5 was ∼115 kDa. Similarly, the molecular mass of FadD5_1-448_ as calculated by SEC-MALLS is ∼96 kDa which is close to the theoretical dimeric molecular mass of ∼101 kDa. Further, the molar mass distribution of the peaks corresponding to FadD5 and FadD5_1-448_ show that they are monodispersed in solution (Fig. S2).

**Fig. 1:**
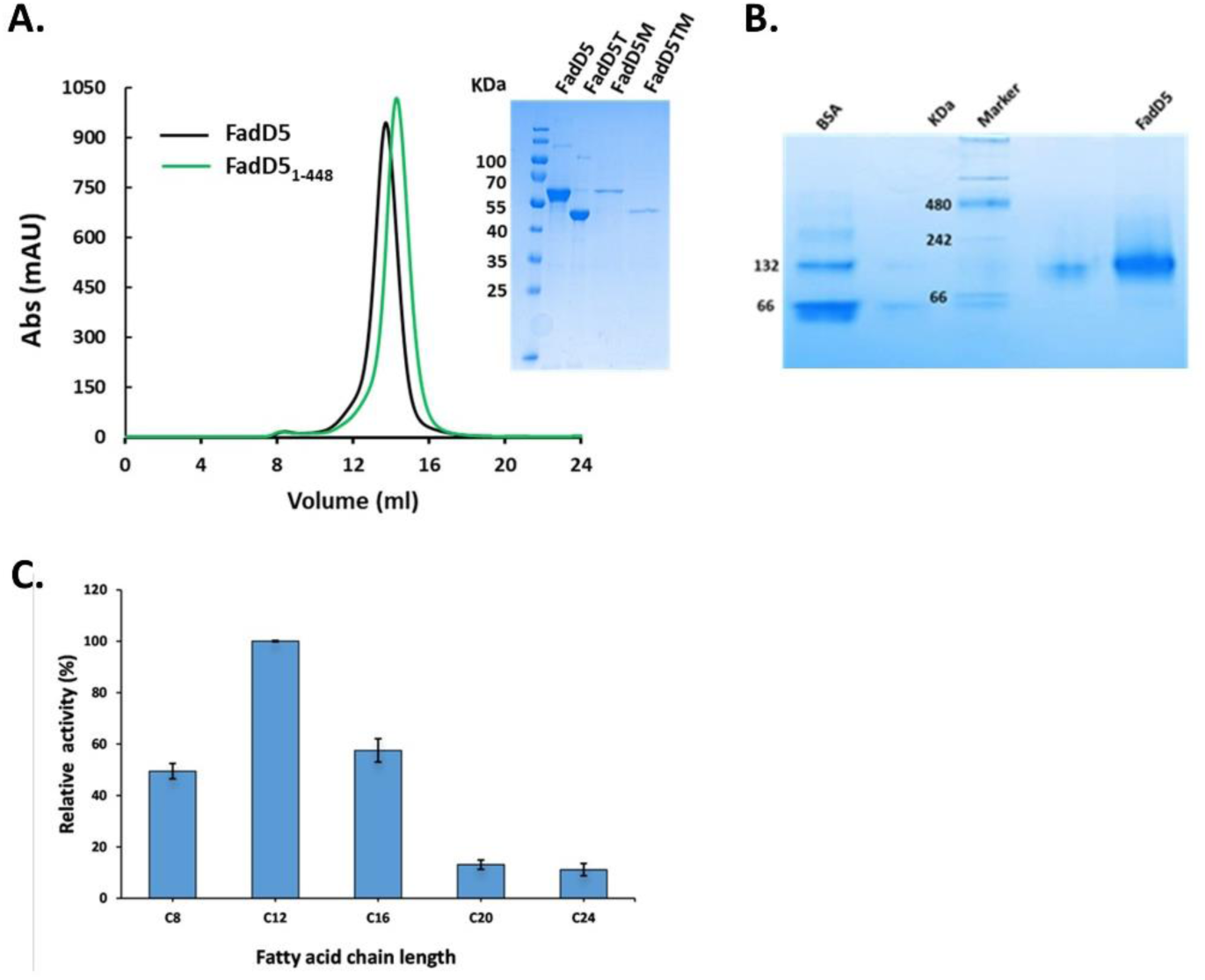
**A**. SEC profile and SDS-PAGE analysis of FadD5 and FadD5_1-448_ from the soluble as well as membrane fraction solubilized using DDM. T=Truncated, M=Membrane fraction. **B.** Blue-native-PAGE analysis for FadD5. **C.** Relative specific activity of FadD5 measured using fatty acids of different chain length from C8 to C24. The activity of FadD5 towards C12 was defined as 100%.

Although we have purified FadD5 as a soluble protein without the use of any detergents, predictions from MemType2 [47] and PPM web server [48] suggested FadD5 to be a peripheral membrane protein. Purification from the membrane fraction of cells expressing FadD5 and analysis with SDS-PAGE clearly showed the presence of FadD5. This confirmed that FadD5 can be associated to membrane although it is stable as a soluble protein. Moreover, a similar study with FadD5_1-448_ variant showed that the N-terminal domain is associated to the membrane (Fig. 1A).

### 3.3 ATP increases the stability of FadD5

CD spectroscopy analysis shows that the purified protein is well folded (Fig. S3A). Melting temperature analysis shows that it has a T_m_ of 43 °C (Fig. S3B). Further, the T_m_ of FadD5 calculated from nanoDSF is 44 °C in close agreement to that calculated by CD. Thermal stability of FadD5 has increased by 3 °C to 47 °C in the presence of ATP. However, the addition of CoA to FadD5 did not change T_m_ (Fig. S4A). The aggregation onset temperature (T_agg_) calculated from the inflection points (IPs)/T_m_ from the nanoDSF data also suggests that ATP has delayed thermal aggregation (Fig. S4B).

### 3.4 FadD5 is a FACS with preference for long chain fatty acids

The sequence analysis already suggested that FadD5 is a FACS with substrate specificity for very long acyl chains. Therefore, enzyme activity of FadD5 was tested with fatty acid substrates with varying acyl chain lengths; C8, C12, C16, C20 and C24 (Fig.1C). While all the tested substrates showed measurable activity, specific activity calculation showed that lauric acid (C12) has the highest specific activity followed by palmitic acid (C16). Further detailed enzyme kinetic studies were performed with C12 and C16 fatty acids. These studies revealed that C16 has a lower K_m_ value compared to C12. However, C12 has higher k_cat_ value than C16 (Table 1). N-terminal domain alone did not show any activity confirming that the C-terminal domain is important to bind the substrate as also demonstrated recently in FadD23 [49].

**Table 1:**
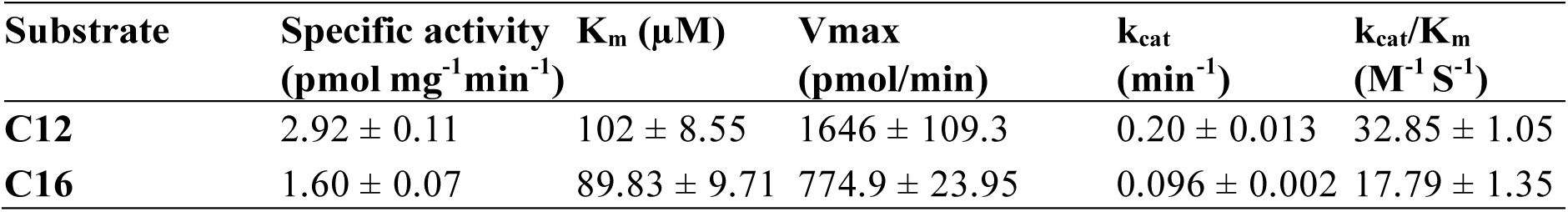
Specific activities and Michaelis-Menten kinetic constants of FadD5 towards C12 and C16 substrate.

To further confirm the FACS activity of FadD5 the formation of the final product was tested through LCMS assays. We have used lauric acid (C12) as the substrate for this test. In the presence of ATP and CoA, FadD5 converts C12 into C12-AMP in the first half of the reaction and then to C12-CoA in the second half of the reaction. The reaction was monitored by following the accurate mass of C12-AMP C_22_H_36_N_5_O_8_P (529.2228 Da) and C12-CoA C_33_H_58_N_7_O_17_P_3_S (949.2765 Da) by selected ion chromatography (SIC). In the control sample, no signal was detected, while the half reaction has a peak for C12-AMP and the full reaction shows both compounds C12-AMP and C12-CoA. The peaks indicated with peak 1 (retention time 5.48 min) corresponds to m/z 528.2228 while peak 2 (retention time 5.7 min) represents m/z 948.2765. The corresponding mass spectra are shown in Fig. S5. Therefore, through enzyme activity assays, we have demonstrated that FadD5 has activity with C12 to C16 fatty acids, providing evidence for its ability to metabolize long chain fatty acids. Tests with C20 and C24 could not be performed further due to solubility issues. It will indeed be interesting to see if FadD5 shows higher catalytic efficiency for these substrates.

### 3.5 FadD5 crystallizes with the C-terminal domain cleaved off

FadD5 was crystallized in sitting drop vapor diffusion method. Interestingly, FadD5 crystals appeared only when the drop contained a particular lot of condition B1 of the Midas plus screen whereas the reservoir solution could be entirely different. This approach is known as ‘alternate reservoir’ method [30]. The crystals obtained diffracted to about 2.8 Å resolution. The crystals belonged to the space group P4_1_2_1_2 with two molecules in the asymmetric unit based on the solvent content analysis. The closest available structural homolog, Mtb FadD13, was used for molecular replacement. Structurally, like other adenylate-forming enzyme family proteins, each monomer of FadD5 is composed of two domains, a larger N-terminal domain (residues 1-427) and a smaller C-terminal domain (residues 434-554) linked by a flexible linker (^428^DRKKDM^433^). Individual domain searches (N– and C-terminal domains) resulted in a solution in which the two N-terminal subunits could be placed with an expected 2-fold symmetry as observed in other FadDs. Despite multiple trials, no solutions were obtained where the C-terminal domain could be placed confidently. However, the N-terminal domain could be built and refined to a final R-free and R of 0.23 and 0.20, respectively. N-terminal 1-19 residues did not show any density, most likely due to the disordered nature of this region. The determined structure of FadD5 is shown in Fig. 2. Crystals obtained were in unliganded form even after incubating the protein with CoA, ATP or CoA+ATP during crystallization. The detailed statistics of data collection and refinement are provided in Table 2.

**Fig. 2:**
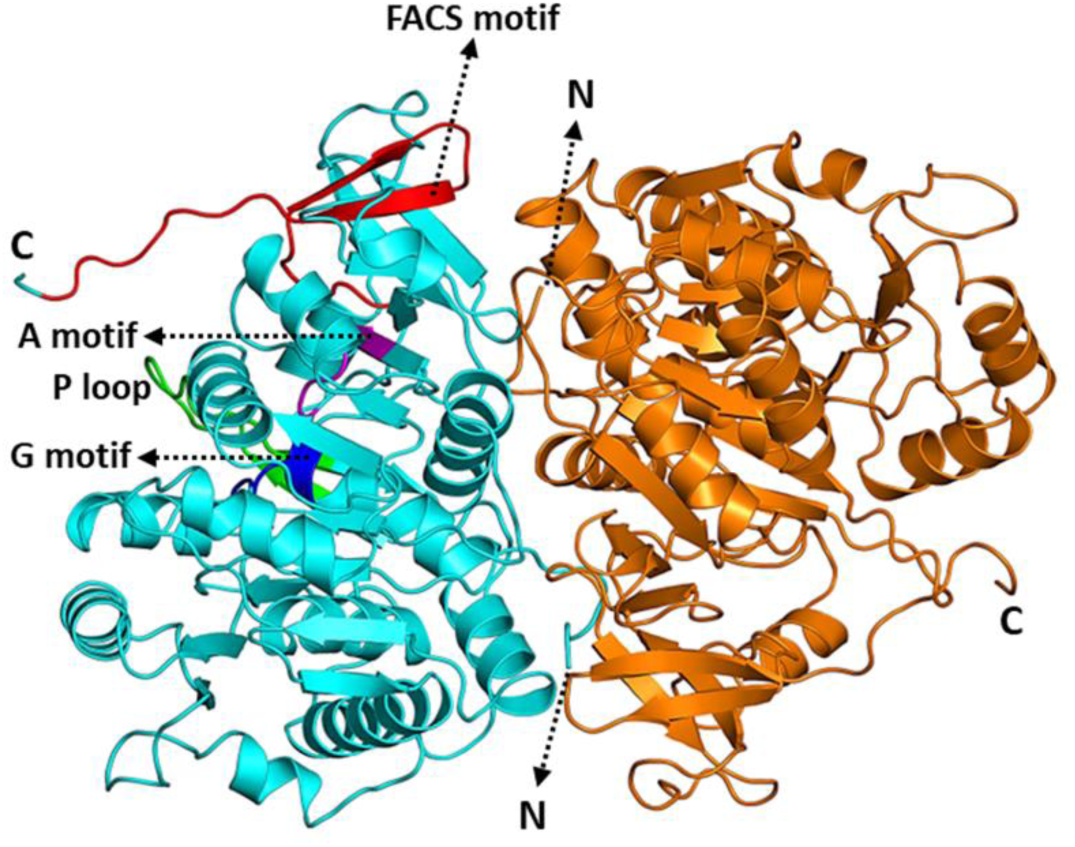
Cartoon representation of the solved crystal structure of FadD5 (amino acids 20-435) with 2 molecules in the asymmetric unit. Chain A in cyan and Chain B in orange. FACS motif in red, A motif in magenta, P loop in green, G motif in blue.

**Table 2:** Data collection and refinement statistics of FadD5.

|  |  |  |
| --- | --- | --- |
| Dataset | FadD5 | FadD5<sub>1-448</sub> |
| Crystallization |  |  |
| Protein Buffer | 20 mM HEPES, pH-7.5; 500 mM NaCl,1 mM BME |  |
| Crystallization solution | 0.1 M Sodium formate, 20 % w/v SOKALAN CP 45 |  |
| Protein concentration | 18 mg ml<sup>-1</sup> |  |
| Cryo protectant | 25% glycerol + Crystallization solution |  |
| Temperature (°K) | 295 |  |
| Data Collection |  |  |
| Beamline | Diamond Light source Beamline I04 |  |
| Detector | DECTRIS EIGER X 16M |  |
| Wavelength (Å) | 0.9795 |  |
| Temperature (°K) | 100 |  |
| Data Processing |  |  |
| Space group | P 4<sub>1</sub> 2<sub>1</sub> 2 | P 4<sub>1</sub> 2<sub>1</sub> 2 |
| Unit cell parameters |  |  |
| a, b, c (Å) | 120.91 120.91 247.91 | 121.67 121.67 249.87 |
| α, β, γ (°) | 90 90 90 | 90 90 90 |
| Processing software | XDS, AIMLESS | xia 2, AIMLESS |
| Resolution range (Å) | 86.56–2.80 (2.90–2.80) | 60.84–2.47 (2.56–2.47) |
| CC<sub>1/2</sub> | 0.99 (0.39) | 0.99 (0.33) |
| R<sub>merge</sub> (%) | 0.13 (2.32) | 0.03 (0.97) |
| R<sub>meas</sub> (%) | 0.14 (2.48) | 0.04 (1.38) |
| R<sub>pim</sub> (%) | 0.05 (0.86) | 0.03 (0.97) |
| I/sigma I | 10.7 (1.0) | 12.5 (0.3) |
| Completeness (%) | 99.7 (100.0) | 97.88 (84.03) |
| Multiplicity | 7.5 (7.8) | 1.9 (1.9) |
| No of reflections | 344933 (34862) | 128204 (8654) |
| No of unique reflections | 45898 (4454) | 67795 (4491) |
| Wilson B-factor (Å<sup>2</sup>) | 79.89 | 73.04 |
| Refinement Statistics |  |  |
| Resolution | 86.56–2.80 | 60.84–2.47 |
| Rwork (%) | 20.52 | 19.79 |
| Rfree (%) | 23.84 | 22.24 |
| No of Atoms | 6331 | 6255 |
| Proteins | 6248 | 6196 |
| Water | 53 | 35 |
| Ligands | 30 | 24 |
| Average B factor ($\text{\AA}^2$) | | |
| Macromolecules | 76.96 | 80.91 |
| Ligands | 79.90 | 100.31 |
| Water | 63.73 | 73.99 |
| RMS deviation |  |  |
| RMS bond ($\text{\AA}$) | 0.002 | 0.003 |
| RMS angle ($^\circ$) | 0.49 | 0.64 |
| Ramachandran plot (%) |  |  |
| Favoured | 95.42 | 97.79 |
| allowed | 4.46 | 2.21 |
| outliers | 0.12 | 0 |
| **PDB id** | 8R2Q | 8R2R |

To understand why the C-terminal domain is missing, we further designed a construct for only the N-terminal domain (FadD5_1-448_; as described in the previous sections). Interestingly, FadD5_1-448_ crystallized under the same conditions, same space group and with the same crystal packing (Table 2). Furthermore, analysis of the crystal packing of the N-terminal domains suggested that there is no space available for the C-terminal domain to be placed in the asymmetric unit/unit cell (Fig. S6). This strongly suggests that the full length FadD5 undergoes proteolytic cleavage for reasons not yet known and the resulting N-terminal domain is crystalized. Noteworthy, the crystals do appear within 3 days. Therefore, we believe that the cleavage of C-terminal domain is mainly a crystallization artefact. Interestingly, the cleavage of C-terminal domain during crystallization is also reported for other Mtb FadDs, namely, FadD13 and FadD28 [4,50].

FadD5 dimer interface is through their N-terminal domain with an interface area of 1818 Å^2^. A total of 19 amino acids from each monomer are involved to form the dimer through intermolecular hydrogen bond formation. The dimeric interface complex formation significance (CSS) score of 0.725 and the solvation energy gain on complex formation (ΔiG) is calculated to be –20.2 kcal/mol. The overall fold of N terminal domain of FadD5 is conserved as in other FACS. The solved crystal structure of FadD5 (N-terminal domain; residues 20-435) superposes well with the N-terminal domain of FadD13 (PDB id: 3r44), *T. thermophilus* FACS (TtLC-FACS; PDB id: 1ult), *N. tabacum* (Nt4CL2; PDB id: 5bsm) with RMSD values of 2.0, 2.3 and 2.3 Å, respectively, as calculated by the SSM superpose [51] option in *COOT* [37].

### 3.6 C-terminal domain movement and SAXS studies on FadD5

One limitation of this study is that we could not get the structure of full-length FadD5. Although the overall fold of N– and C-terminal domains are conserved between various FACS, the relative position of C-terminal domain with respect to N-terminal domain is very different in different FACS (Fig. S7). The N– and C-terminal domains are linked by a flexible linker and the C-terminal domain moves as a rigid body and takes different conformation either in acyl-adenylate/open conformation or acyl-CoA-forming (thio)/closed conformation with respect to the N-terminal domain based on the bound ligand [40,52,53]. For example, in plant 4-coumarate CoA ligase (Nt4CL2) C-terminal domain rotates as much as 140° from the adenylate conformation upon binding to CoA [8]. It happens due to the displacement of core motif A10 away from the active site and exposed to solvents, leading to a new set of residues stabilizing the bound nucleotide. Additionally, this change was accompanied by a more closed P loop structure facilitated by the insertion of the A8 loop [53]. In the absence of the full-length crystal structure of FadD5, we utilized *in silico* methods to reconstruct the C-terminal domain and generate the full-length FadD5 structure. The predicted model from AlphaFold server [54] was already in adenylate conformation. Monomeric AlphaFold predicted structure was superposed to the crystal structure dimer to generate full length FadD5 dimer (FadD5_AF_) in adenylate conformation. In addition, the AlphaFold predicted structure of C-terminal domain (residues 430-554) was superposed on other FACS enzyme TtLC-FACS (PDB id: 1ult, 1v25 and 1v26), FadD13 (PDB id: 3r44), and NtCL2 (PDB id: 5bsr) to generate full length FadD5 models with different C-terminal domain conformations, which were refer here as FadD5_1ult_, FadD5_1v25_, Fad5_1v26_, FadD5_3r44_ and FadD5_5bsr_, based on the reference PDB id. In Fig. 3 we show two conformations of C-terminal domain to highlight the possible C-terminal domain movement (Fig. 3) [53].

**Fig. 3:**
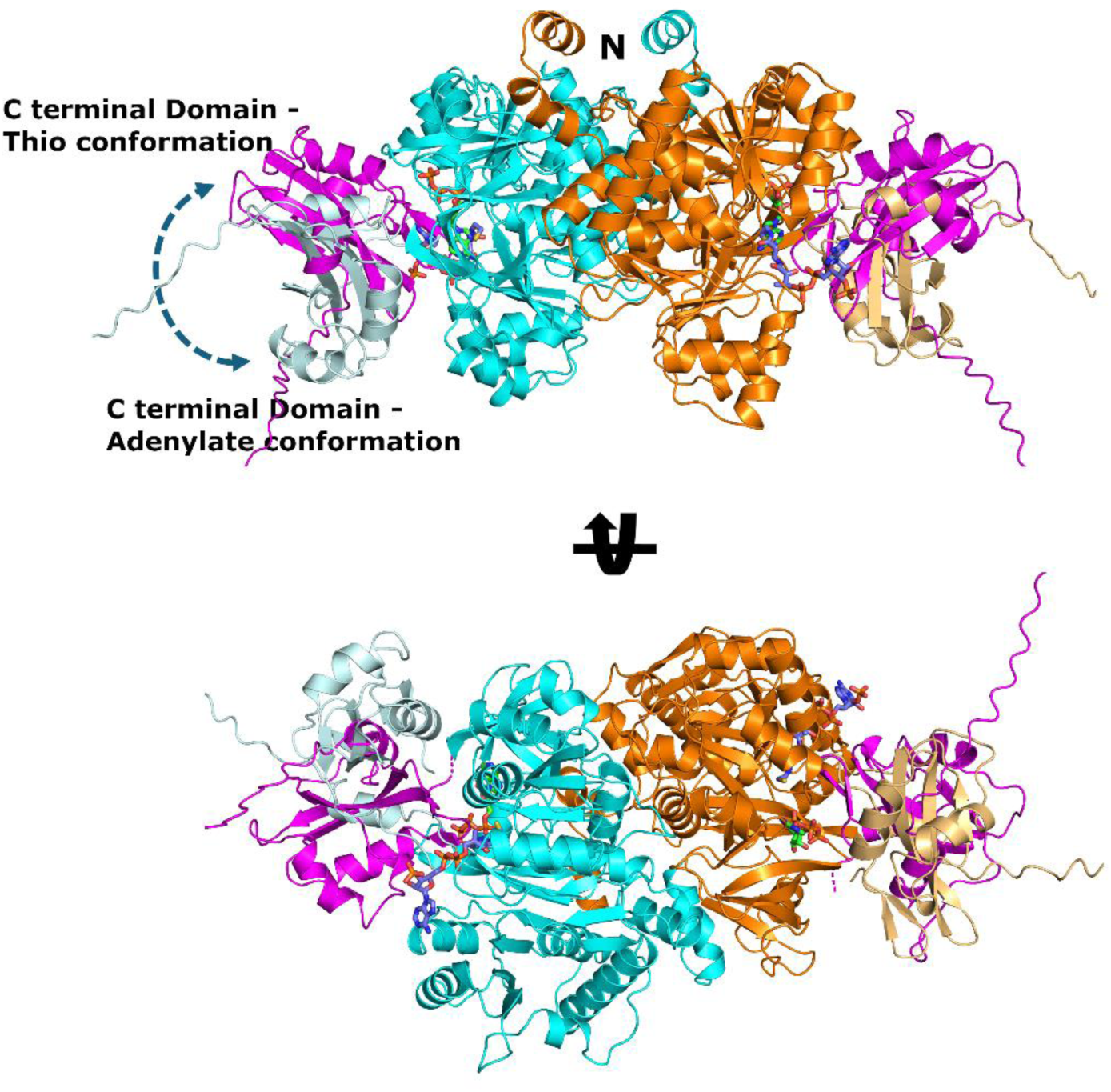
Cartoon representation of the modelled FadD5_AF_ and FadD5_5bsr_. The two N-terminal domains are shown in cyan and orange. The conformation of C-terminal domain as in FadD5_AF_ and FadD5_5bsr_ are shown in light cyan and magenta cartoon, respectively. Modelled ATP and CoA based on structural superposition is shown in green and blue sticks, respectively, showing the possible binding sites in FadD5.

In addition to the missing C-terminal domain in the crystal structure, obtaining ligand-bound complex structures through co-crystallization or soaking were unsuccessful, despite extensive efforts, most likely due to the absence of the C-terminal domain in the crystals. Therefore, to further understand whether FadD5 stays as a full-length stable dimer in solution and to identify if there are any C-terminal domain movements due to the binding of ligands, we performed SAXS studies on FadD5 in the unliganded form as well as in the presence of ATP or CoA. The dimensionless Kratky plot analysis suggests that FadD5 is well folded (Fig. 4) with an *R_g_* of 35.5 Å and *D_max_* of 123 Å. The calculated molecular weights were close to the expected molecular mass of FadD5 dimer (Table 3). Different models of FadD5 were fitted against these SAXS scattering profiles using FoXS [55]. The solved structure of FadD5 containing only the dimer of N-terminal domain or the full-length monomer (AlphaFold predicted structure) does not fit very well with the experimental curve with the respective *χ^2^* values of 39 and 87. However, full length FadD5 dimer models fitted well to the SAXS data (Table 3) suggesting that FadD5 exists in its full-length and dimeric form further confirming that C-terminal cleavage is only a crystallization artefact. Among the tested models, FadD5_AF_, FadD5_3r44_, and FadD5_1ult_ fitted better while the other models showed a downward bump near 0.13 q Å^-1^. It should be noted that each of these good fitting models still have C-terminal domain in slightly different conformations. The *R*_g_, *D*_max_ and the molecular mass calculated for FadD5 in the presence of ATP or CoA were very similar to that of the unliganded FadD5 (Table 3). Also, as in the case of unliganded form, FadD5 dimer models based on FadD5_AF_, FadD5_3r44_ and FadD5_1ult_ fitted better to the SAXS data of FadD5 in the presence of ATP. For FadD5 SAXS data collected in the presence of CoA, the fit was better for the model with C-domain conformation as in FadD5_1ult_. Even for this fit, the C2 value as defined in FoXS [55] is 3.9. For other models, the C2 values were 4 or higher, indicating overfitting. It is possible that the conformation for FadD5 C-terminal domain is different from those in the tested models in the presence of CoA. Higher resolution data is needed to ascertain this.

**Fig. 4:**
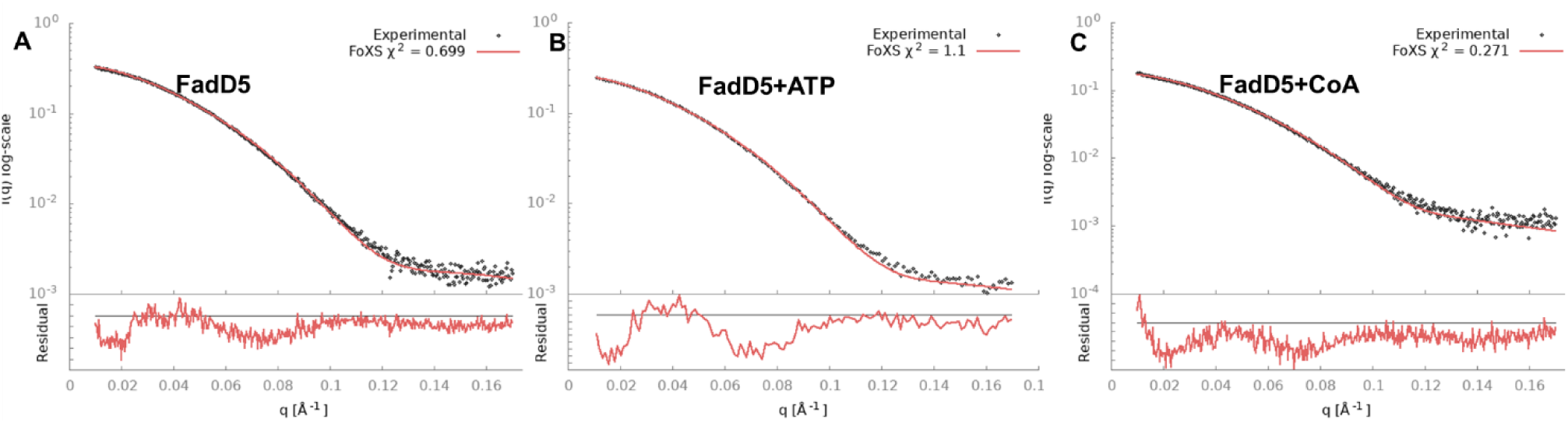
SAXS curves for FadD5 collected in **A.** unliganded form **B.** in the presence of ATP and **C.** in the presence of CoA. The fit of the structural models (**A** & **B.** FadD5_AF_; **C.** FadD5_1ult_) to the experimental scattering curve is shown in red. The residuals are plotted at the bottom of the curve.

**Table 3.**
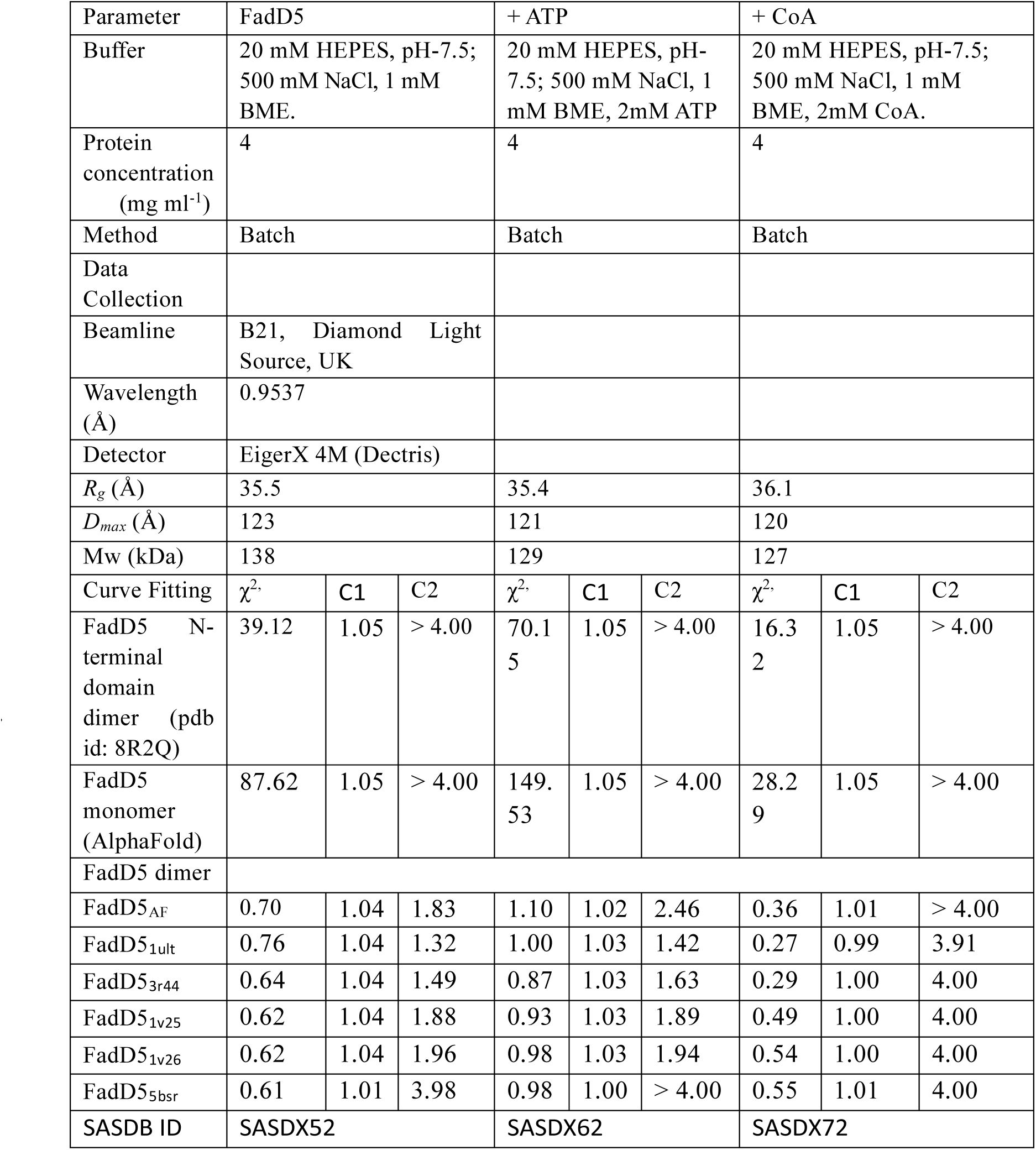
*R_g_* and *D_max_* for Apo-, ATP– and CoA-bound FadD5 were calculated from ScÅtter program, and the molecular mass was estimated with the Porod invariant (Qp) method.

### 3.7 ATP and CoA binding sites of FadD5

For the modelling studies, FadD5_AF_ was used. ATP molecule was modelled by the superposition of ATP bound structure of Nt4CL2 (PDB id: 5bsm) [53] onto FadD5_AF_ (Fig. 5) and residues within 5 Å radius of the modelled ATP have been identified to understand important residues interacting with ATP molecule (Fig. 5E). When compared, we found that the adenosine molecule is enveloped by the core motif A5/A motif, denoted as ^330^AAFGQTE^336^. Notably, Glu336 is highly conserved, but variations are observed in Ala330, where Gln299 is present in FadD13 and Gln331 in Nt4CL2. Additionally, Phe332 is substituted by Tyr301 and Tyr333 in FadD13 and Nt4CL2, respectively. The P-loop, also known as motif A3, engages with the ATP molecule through the highly conserved sequence ^188^IMYTSGTTGRPKGA^201^. Lys522, situated in core motif A10 and being the sole residue from the C-terminal domain, plays a crucial role in the adenylation reaction [53]. Mutations affecting His252 and Thr351, corresponding to His236 (core motif A4) and Thr335 (core motif A5), have been shown to impair activity in previous studies [53]. CoA molecule was modelled on to FadD5_AF_ by the superposition of CoA bound structure of Nt4CL2 (PDB id: 5bsr) [53]. Residues ^231^GVPXXHIAG^239^ belong to the G motif. In this conformation, there were no contacts between CoA and the C-terminal domain residues (Fig. 5). Comparison of residues interacting with CoA in FadD5_AF_ and Nt4CL2 suggests that FadD5 has smaller residues. For example, residues Gly231, Val232, Ala238, Gly239, Ala260, Gly437, and Glu439 in FadD5 correspond to Val232, Leu233, Tyr239, Ser240, Lys260, Lys443, and Phe445 in Nt4CL2. Residues within a 5 Å radius of modelled CoA in FadD5_AF_ are shown in Fig. 5E.

**Fig. 5:**
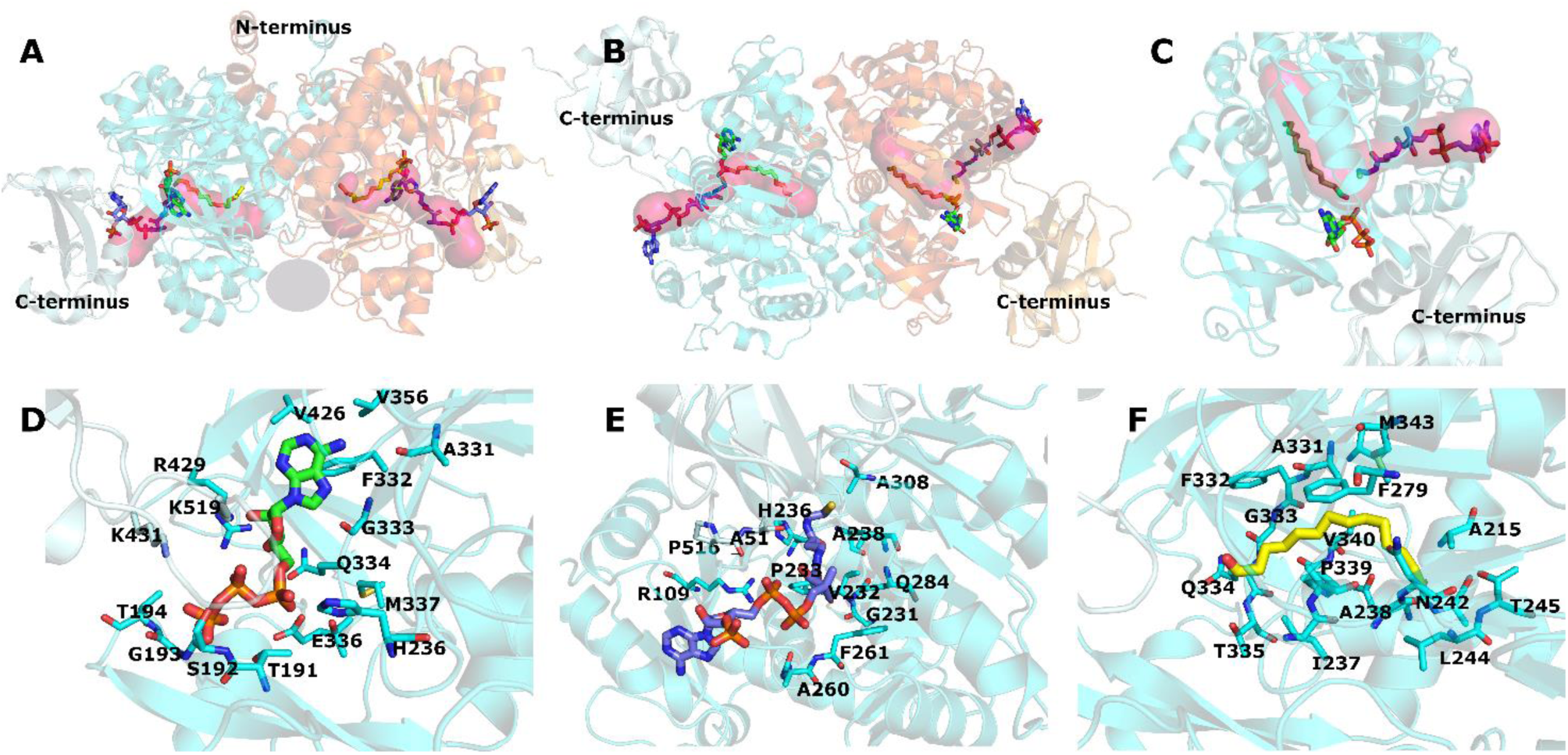
Ligand modelling studies on FadD5_AF_. A &B. Predicted binding pocket for fatty acyl tail based on structural superposition with PDB id 1v26. The predicted binding pocket from Mol2 analysis [56] is shown in surface representation in magenta. CoA, ATP and myristic acid are shown in blue, green and yellow sticks. The space between the two monomers is highlighted in grey through which longer acyl tails can be accommodated. C. Monomer of FadD5 showing the fatty acyl binding pocket with bound ATP, COA and myristic acid. D-E. Residues that are within 5 Å radius of modelled ATP (D.), COA(E.) and MYR(F.)

### 3.8 Fatty acyl tail binding pocket of FadD5

In TtLC-FACS, Trp234 has been described as a gating residue that opens the fatty acid binding tunnel followed by ATP binding. However, the corresponding Tyr234 is replaced with Ile240 in FadD5 suggesting that no gating is required for the binding of long chain substrates to FadD5. Further comparison of FadD5_AF_ with the myristoyl-AMP bound structure of TtLC-FACS (PDB id: 1v26) [40] reveals a putative fatty acid binding pocket in FadD5 (Fig. 5A, 5B, 5C). Structural analysis suggests that Ile240 in FadD5 is pointing away from the proposed fatty acid binding tunnel. Analysis of the putative fatty acid binding pocket suggests that it could accommodate substrates only slightly longer than C14. The predicted acyl binding pocket is approaching the dimer interface and therefore may interfere with the binding of very long acyl chains (Fig. 5A, 5B). However, our enzyme activity studies shows that FadD5, can also activate fatty acids longer than C16. FadD13, which is also known to work with very long fatty acyl chains and the closest homolog of FadD5 has been shown to be in an equilibrium of monomer and dimer. It has been proposed that the dimeric form of FadD13 stays in the solution while and the monomer binds to the membrane. A possible membrane binding region has been identified in FadD13. Membrane association of FadD13 through this region allows the binding of very long chain fatty acids [11]. A comparison of the predicted membrane binding region of FadD13 (Fig. 6A) with the corresponding region of FadD5 (Fig. 6B) suggests that the surface properties of FadD5 are different. As FadD5 is associated with *mce1* operon and likely processes the lipids imported through the Mce1 complex, it might act differently when compared to FadD13 for membrane association. In another FACS, TtLC-FACS, the longer tails are proposed to be accommodated through a tunnel facing the space between the two monomers. The corresponding space in FadD5 is highlighted with a grey ellipse in Fig. 5A. FadD5 has an extended N-terminal region which is disordered in the crystal structure. However, structure prediction from AlphaFold [54] suggests this region to be composed of a helix with a few positively charged residues and predominantly hydrophobic surface (Fig. 6C). It is possible that this region may have a role in membrane association. Indeed, our studies also show that the N-terminal domain of FadD5 has the ability to associate to membrane. Therefore, we propose that the long chain fatty acids imported by the Mce1 complex to the cytoplasm are transferred to FadD5 through the space between the two monomers while FadD5 is attached to the membrane via the N-terminal helix. This is schematically shown in Fig. 7.

**Fig. 6:**
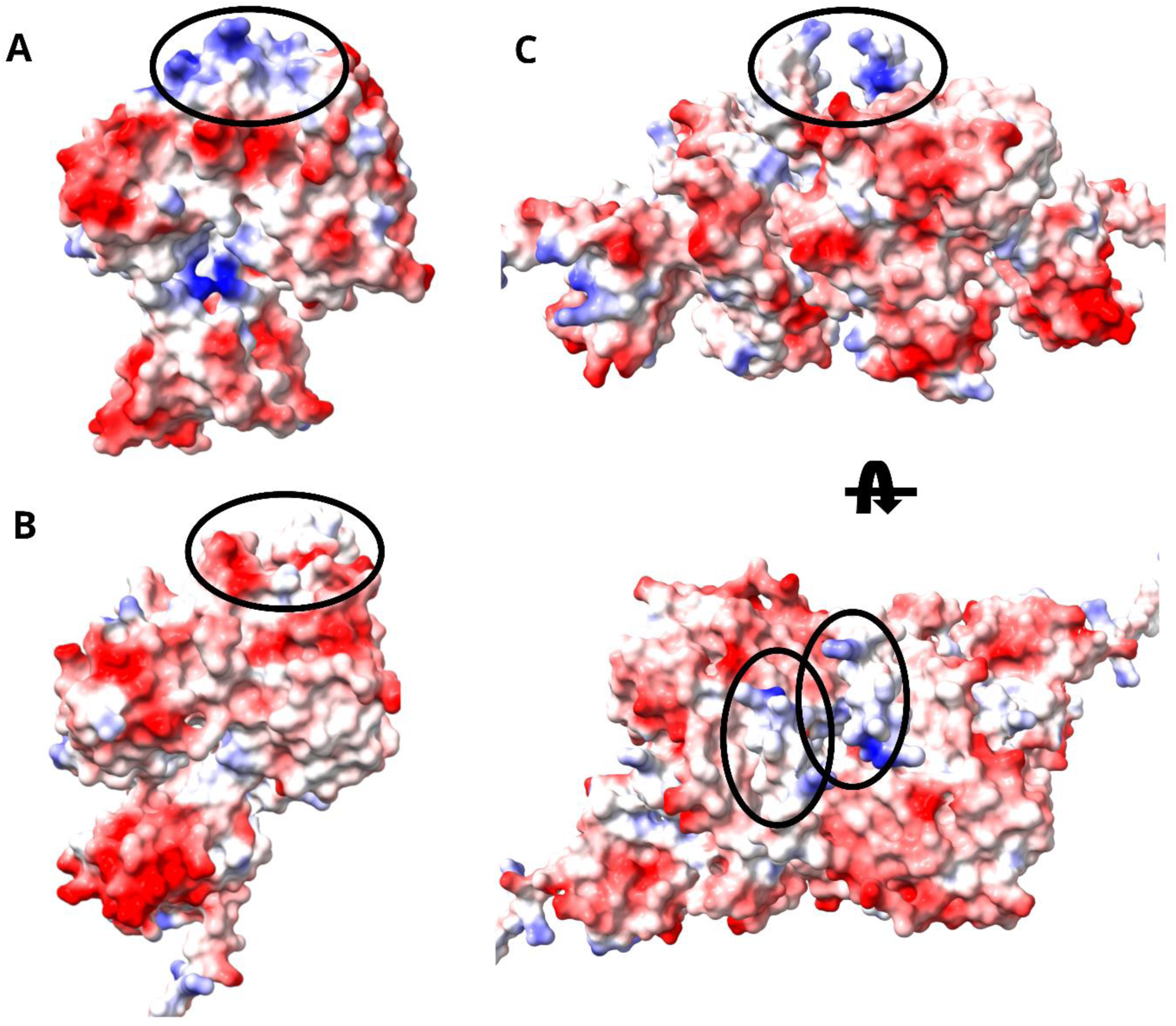
Electrostatic surface analysis of Membrane binding properties of FadD5 as compared to FadD13. A. Electrostatic surface potential of FadD5_AF_ in two orientations. The N-terminal helices are highlighted in black oval in both the orientations. B. Electrostatic surface potential for FadD13 monomer. The potential membrane binding site as proposed by Lundgren et al [57] is highlighted in black oval. C. Electrostatic surface potential of FadD5 monomer. The surface corresponding to that highlighted in B. is highlighted in black oval showing different charge distribution in FadD5.

**Fig. 7:**
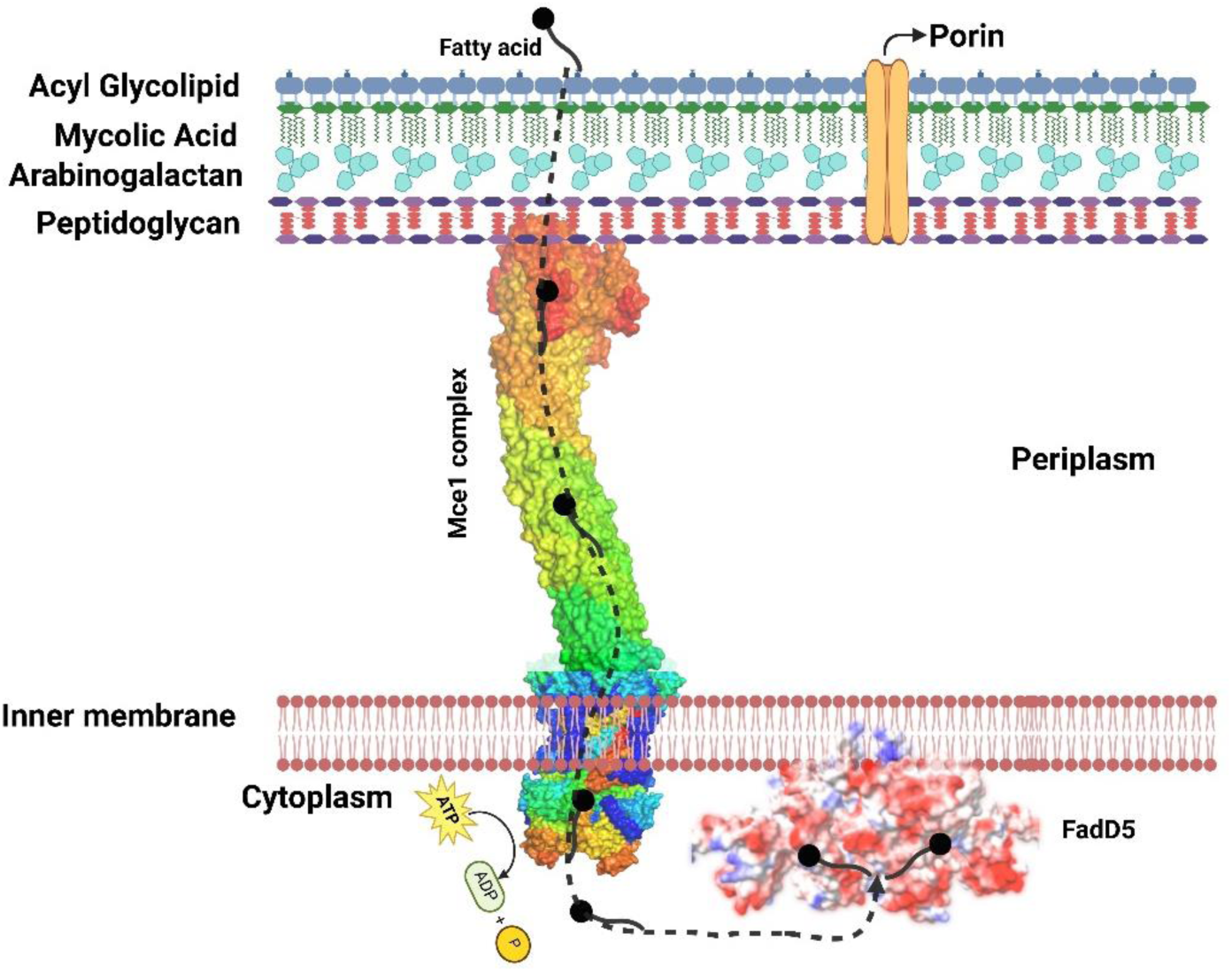
A schematic representation of fatty acid transfer to FadD5. The structure of *M. smegmatis* Mce1 complex [17] is shown in surface representation as embedded to the Mtb cell envelop. FadD5_AF_ represented in electrostatic surface potential is shown as associated to the membrane via the N-terminal helices. Fatty acids imported by the Mce1 complex can bind to the FadD5 via the space between the two monomers as schematically shown by dotted arrows.

## 4. Concluding remarks

Mtb demonstrates remarkable metabolic versatility due to its unique fatty acid metabolism system. It possesses a large amount of fatty acid metabolism genes, including FACSs and FAALs, pivotal in activating fatty acids for metabolic pathways. FadD5, a putative FACS is encoded in the *mce1* operon in Mtb. Mce1 complex encoded by the *mce1* operon is a mycolic and fatty acid transporter. FadD5 being a FACS will have a central role in activating the fatty acids imported through the Mce1 complex for further processing by pathways such as β-oxidation. In this study we demonstrated the FACS activity of FadD5 in the conversion of fatty acids to fatty acyl-CoAs. ATP and CoA binding influence FadD5’s stability and conformation. FadD5 has a dimeric arrangement in its active soluble form and the C-terminal domain is crucial for its activity. This study on FadD5 suggests that it likely acts as a peripheral membrane protein, which is crucial for anchoring longer fatty acyl substrates. Although the full-length crystal structure and the structural flexibility between N– and C-terminal domains still remain unknown, modelling and solution structure studies suggests that, possibly, the extended N-terminal region of FadD5 will have role in membrane anchoring allowing very long fatty acyl chains to bind through the space between the two monomers and thus provides insights on to the possible mode of fatty acyl tail binding.

## Data availability statement

All data generated in this study are included in this article.

## Supplementary Material

Supplementary data is included as a separate file:

## CRediT authorship contribution statement: Mohammad Asadur Rahman

Writing–original draft, Writing – review & editing, Conceptualization, Methodology, Formal analysis, Investigation, Visualization. **Subhadra Dalwani:** Writing–review & editing, Methodology, Formal analysis, Visualization. **Rajaram Venkatesan**: Conceptualization, Writing-original draft, Writing–review & editing, Investigation, Methodology, Supervision, Funding acquisition, Formal analysis, Visualization.

## Supporting information

MtbFadD5_SupplementaryFigures

## Abbreviations

Mtb: Mycobacterium tuberculosis
MCE: mammalian cell entry
FACS: Fatty acyl-CoA synthetase
FAAL: fatty acyl-AMP ligase

## Acknowledgements

We acknowledge and cherish the memory of Prof Lee Riley (Deceased) from the University of California, Berkley, USA for his advice and support. We acknowledge the support from Diamond Light Source for X-ray and SAXS data collection. We acknowledge Prof. Jan Skov Pedersen, Aarhus University for help in rebinning the SAXS data. We also acknowledge the support from Biocenter Oulu structural biology, proteomics and protein analysis, and DNA sequencing core facilities.

## Ethical statement

Not applicable.

## Consent for publication

Not applicable.

