## Supplementary material for "Structural enzymological studies of the long chain fatty acyl-CoA synthetase FadD5 from the *mce1* operon of *Mycobacterium tuberculosis*": MtbFadD5_SupplementaryFigures

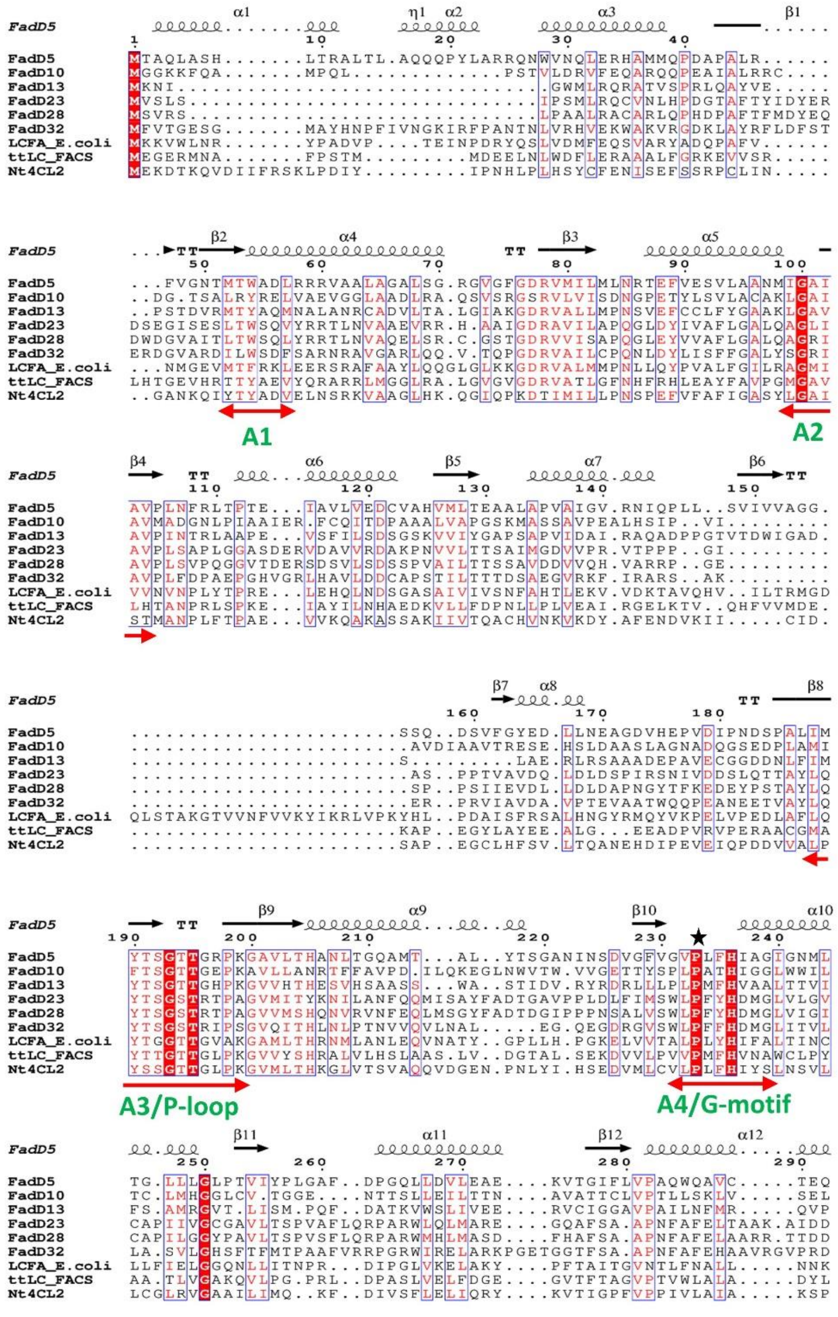
A1
A2
A3/P-loop
A4/G-motif

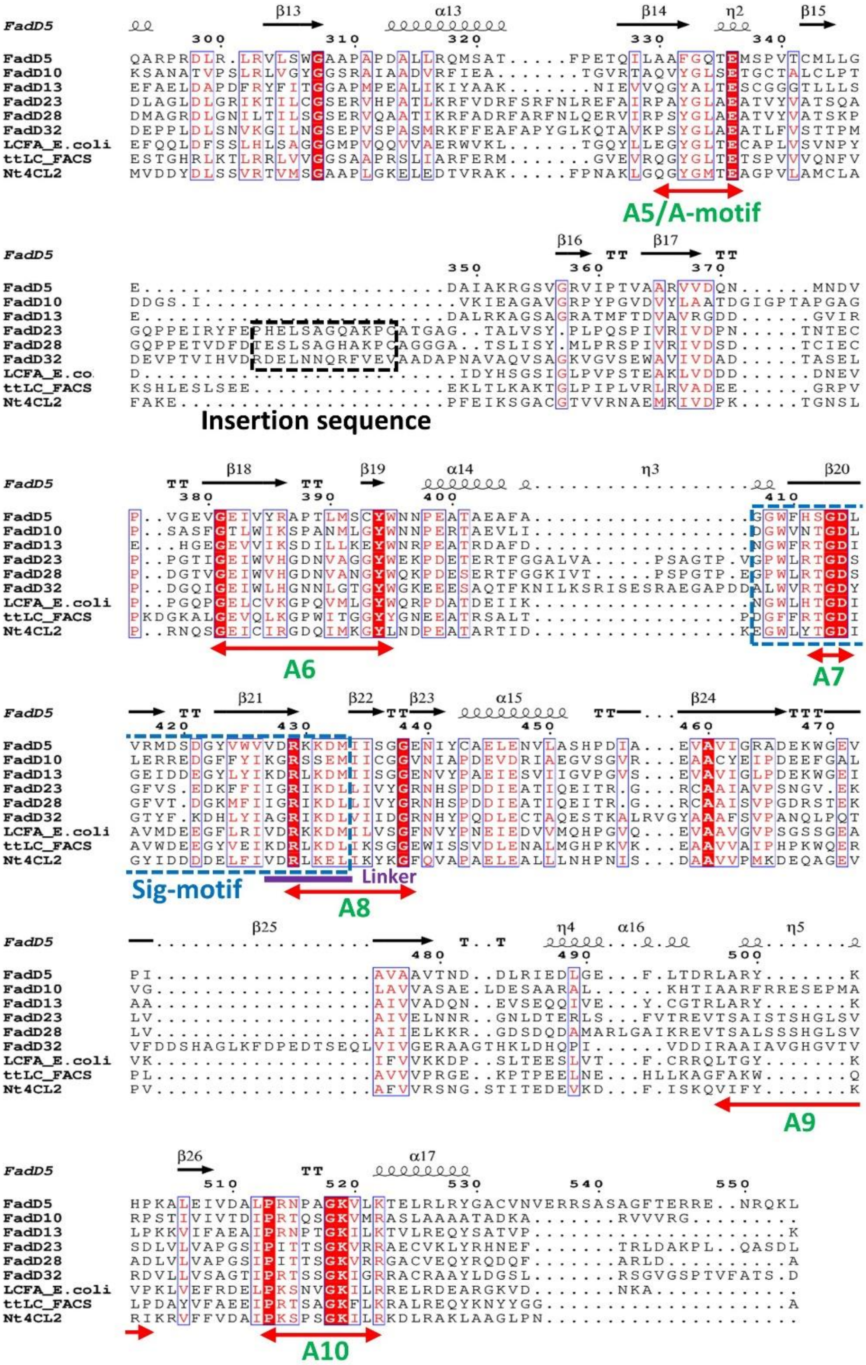


**Fig S1:** Multiple sequence alignment of FadD5. Sequence alignment was generated using T-Coffee program on EMBL-EBI website [1] and then visualized using ESPRIPT server [2]. The secondary structure is from the predicted structure of FadD5 by AlphaFold2 [3]. Important domains and motifs are highlighted with double headed arrows below the sequences.

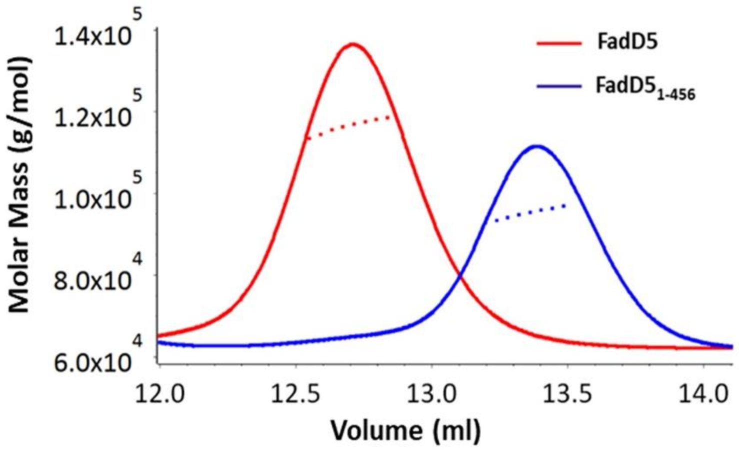


**Fig S2:** The SEC-MALLS profile of FadD5 and FadD5$_{1\text{-}456}$ on a 24 ml Superdex 200 10/300 increase column. The X-axis shows the elution volume and Y-axis shows the molar mass distribution (g/mol). MALLS signal for full length FadD5 is shown in red lines and moler mass distribution in red dotted lines. On the other hand, MALLS signal for FadD5$_{1\text{-}456}$ is shown in blue lines and molar mass distribution in blue dotted lines.

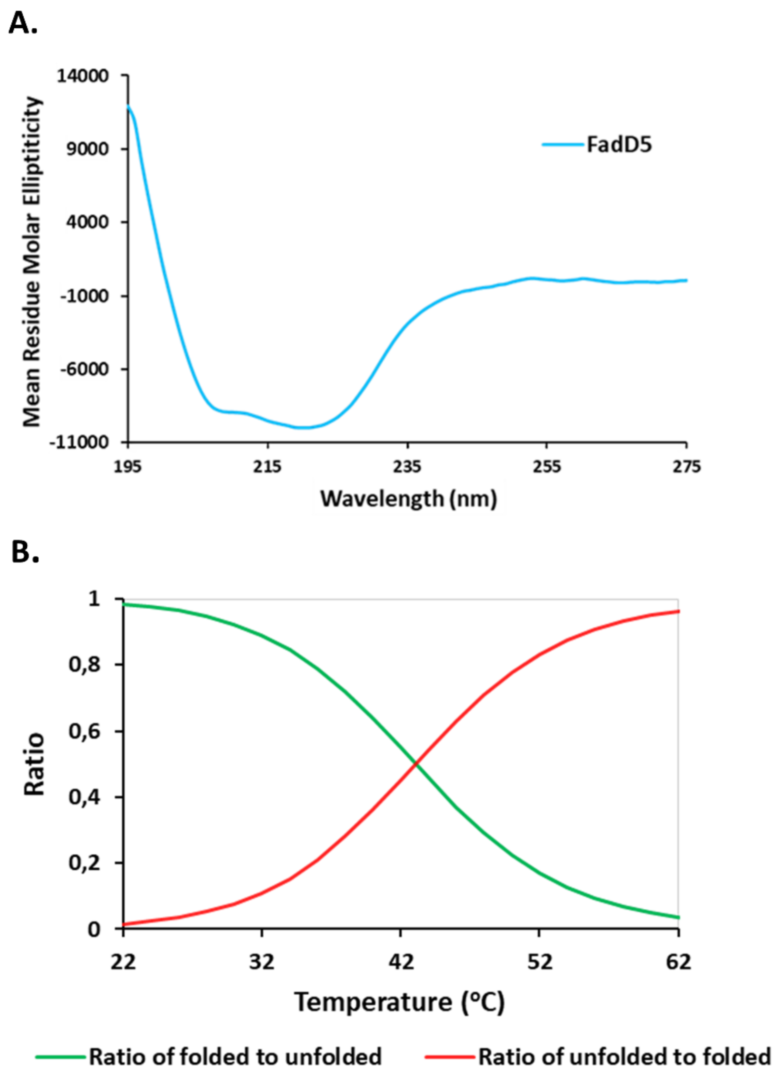


**Fig S3:** CD and thermal stability analysis of FadD5. (A) Graphs showing Far UV CD spectra of FadD5 at 22 °C. (B) Melting curve of FadD5.

**A.**

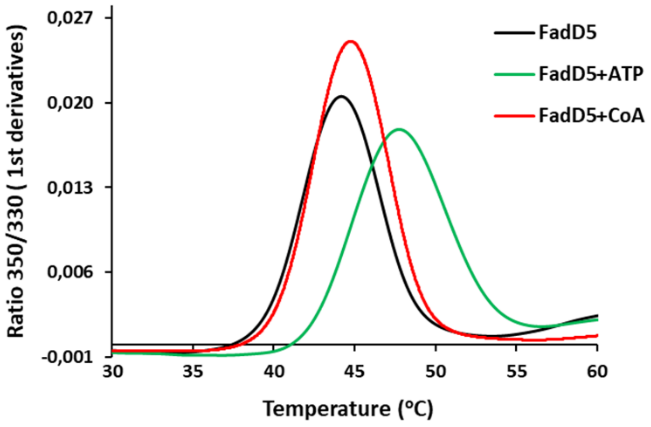


**B.**

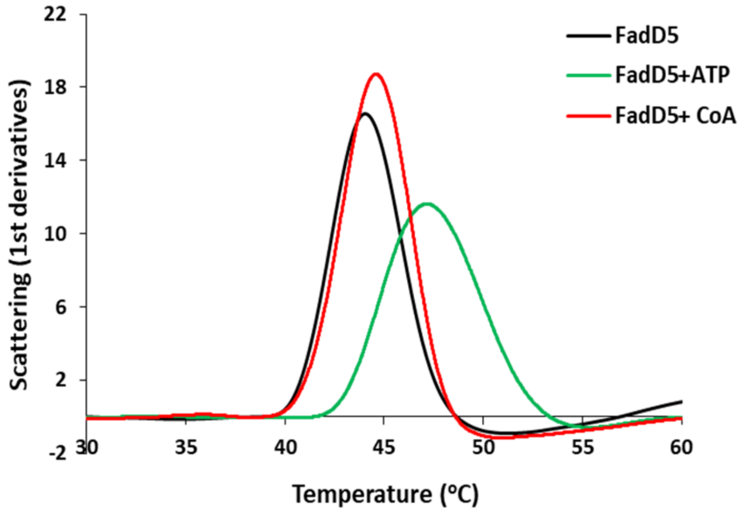


**Fig S4:** Thermal stability analysis of FadD5 with nanoDSF. FadD5 is shown in black, FadD5+2mM CoA in Red and FadD5+2mM ATP in green curves. (A) DSF (B) Scattering.

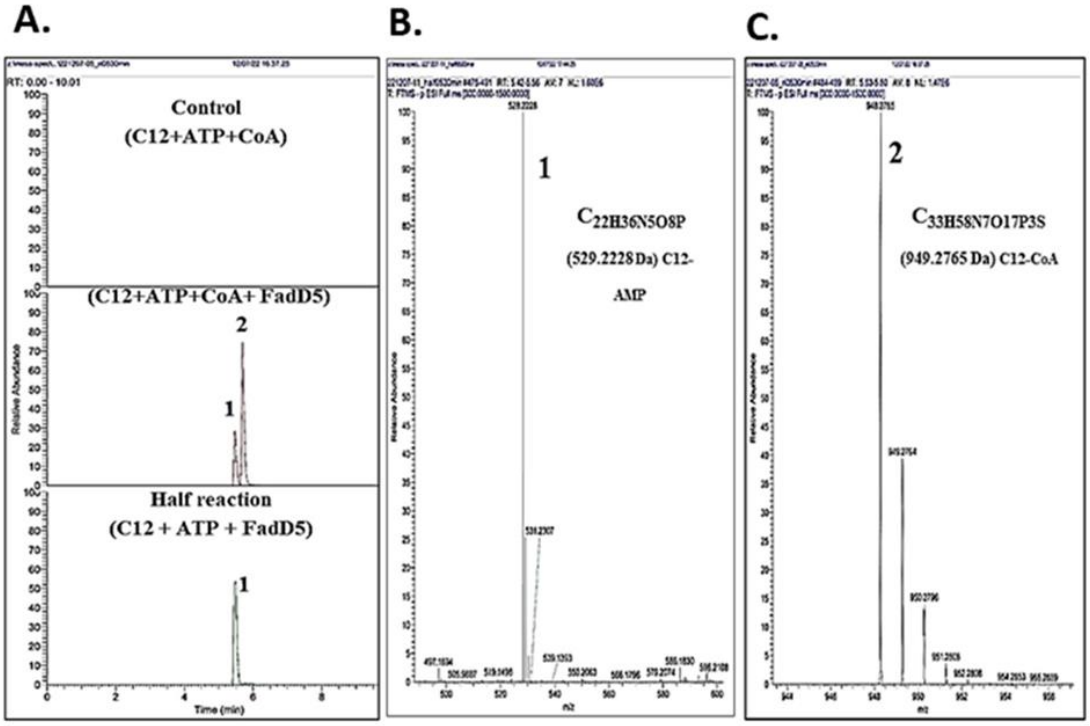


**Fig S5:** (A) FadD5 reaction monitoring and SIC extracted at ± 5 milli mass unit (mmu) for the sum of C12-AMP, peak 1 and C12-CoA peak 2. (B) Mass spectra for the peak 1 and (C) peak 2 in the SIC of the FadD5 reaction.

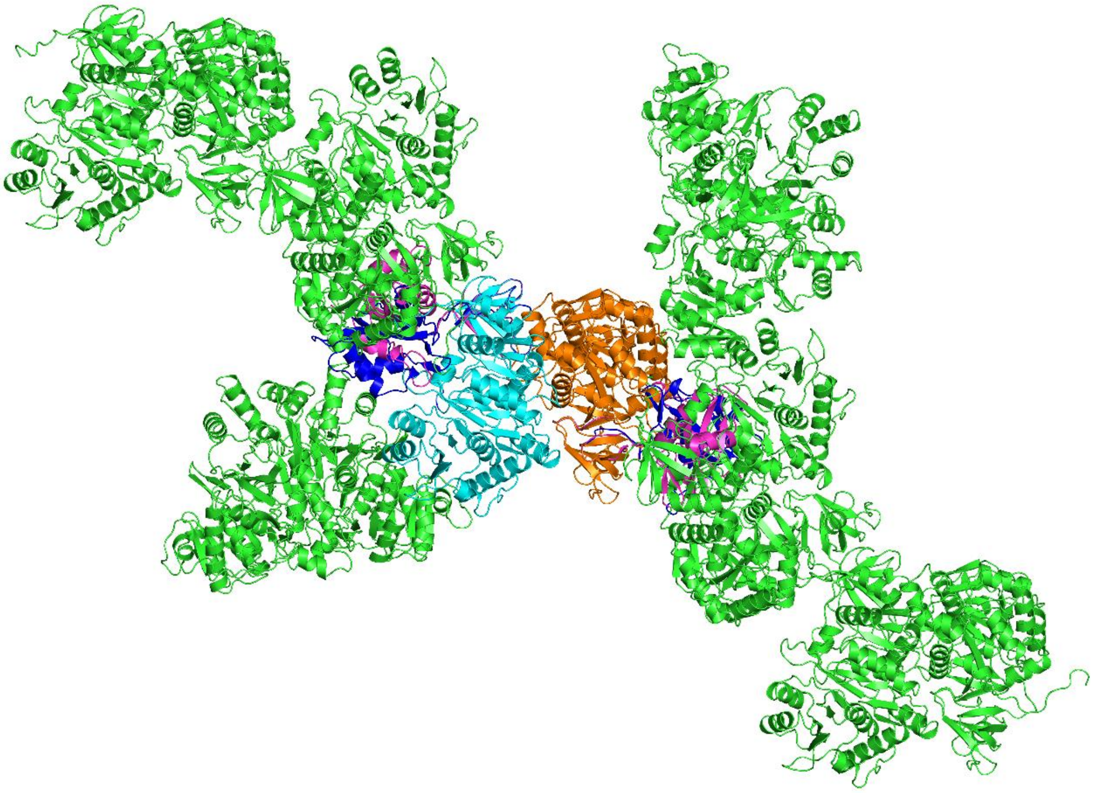

**Fig S6**: Cartoon representation of crystal packing as observed in FadD5 crystals. The two molecules of FadD5 in the asymmetric unit are shown in cyan and orange cartoon. The symmetry related molecules are shown in green cartoon. The missing C-terminal domain modelled in two possible conformations (adenylate conformation in magenta and thio conformation in blue- as discussed later in section 5.10) shows the possible clashes of these domains with the symmetry mates suggesting that the C-terminal domain is not present in the crystals.

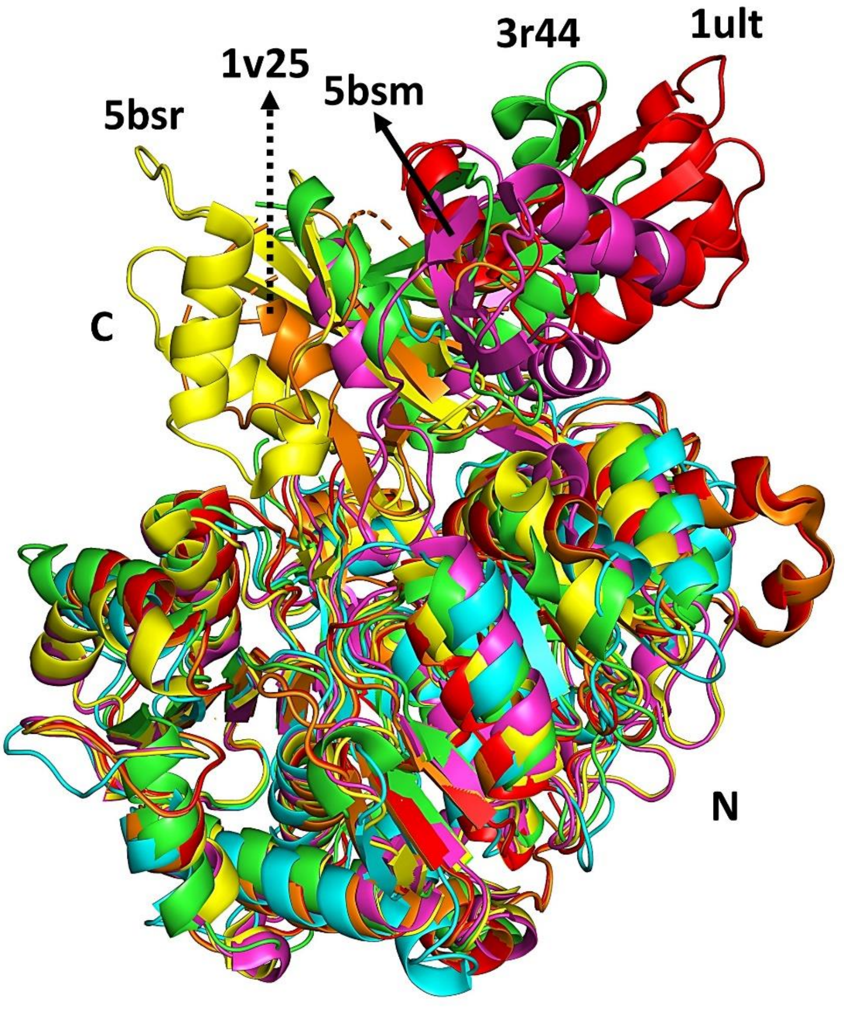


**Fig S7:** Superposition of several structures to show the positioning of the C terminal domain. FadD5-cyan (N-domian); 3R44-green (Apo); TtLCFACS_1ult- red (apo) and 1v26-orange (Myristic acid and AMP bound); Nt4CL2_5bsm-magenta (ATP) and 5bsr- yellow (CoA and AMP).
